# Logic-based mechanistic machine learning on high-content images reveals how drugs differentially regulate cardiac fibroblasts

**DOI:** 10.1101/2023.03.01.530599

**Authors:** Anders R. Nelson, Steven L. Christiansen, Kristen M. Naegle, Jeffrey J. Saucerman

## Abstract

Fibroblasts are essential regulators of extracellular matrix deposition following cardiac injury. These cells exhibit highly plastic responses in phenotype during fibrosis in response to environmental stimuli. Here, we test whether and how candidate anti-fibrotic drugs differentially regulate measures of cardiac fibroblast phenotype, which may help identify treatments for cardiac fibrosis. We conducted a high content microscopy screen of human cardiac fibroblasts treated with 13 clinically relevant drugs in the context of TGFβ and/or IL-1β, measuring phenotype across 137 single-cell features. We used the phenotypic data from our high content imaging to train a logic-based mechanistic machine learning model (LogiMML) for fibroblast signaling. The model predicted how pirfenidone and Src inhibitor WH-4-023 reduce actin filament assembly and actin-myosin stress fiber formation, respectively. Validating the LogiMML model prediction that PI3K partially mediates the effects of Src inhibition, we found that PI3K inhibition reduces actin-myosin stress fiber formation and procollagen I production in human cardiac fibroblasts. In this study, we establish a modeling approach combining the strengths of logic-based network models and regularized regression models, apply this approach to predict mechanisms that mediate the differential effects of drugs on fibroblasts, revealing Src inhibition acting via PI3K as a potential therapy for cardiac fibrosis.

**Significance:** Cardiac fibrosis is a dysregulation of the normal wound healing response, resulting in excessive scarring and cardiac dysfunction. As cardiac fibroblasts primarily regulate this process, we explored how candidate anti-fibrotic drugs alter the fibroblast phenotype. We identify a set of 137 phenotypic features that change in response to drug treatments. Using a new computational modeling approach termed logic-based mechanistic machine learning, we predict how pirfenidone and Src inhibition affect the regulation of the phenotypic features actin filament assembly and actin-myosin stress fiber formation. We also show that inhibition of PI3K reduces actin-myosin stress fiber formation and procollagen I production in human cardiac fibroblasts, supporting a role for PI3K as a mechanism by which Src inhibition may suppress fibrosis.

## Introduction

Cardiac fibroblasts are the primary regulators of remodeling following cardiac injury^1^. Extracellular matrix (ECM) deposition by activated myofibroblasts is essential to this response, but excessive deposition can lead to ventricular stiffness, diastolic dysfunction, and heart failure^1^. While fibroblasts are critical to the wound healing response, current standard-of-care therapeutics for cardiac injury, such as myocardial infarction (MI), affect downstream symptoms but do not specifically target fibroblast signaling^2^. Recent drug discovery and development has focused on identifying drugs such as Entresto (sacubitril/valsartan) that reduce fibrosis in part by modulating fibroblast signaling^3, 4^.

Collagen secretion, αSMA expression, and actin filaments (F-actin) are traditional markers for a profibrotic fibroblast phenotype^5, 6^. While high expression of these markers provides an initial indication of myofibroblast activation, traditional marker expression is inconsistent and does not fully capture the fibrotic response^7^. Recent studies of fibroblast phenotype have shown that fibroblasts exhibit high phenotypic heterogeneity across many facets in response to injury, and that phenotypic changes are also sensitive to drug perturbations^8–11^. Identifying drugs that regulate fibroblast signaling may provide targeted control of fibrosis.

Previously, we developed a logic-based mechanistic network model of fibroblast signaling and applied it to perform virtual screens for anti-fibrotic drugs^12, 13^. That study predicted and experimentally validated an antifibrotic role for the TGFβ receptor inhibitor galunisertib^13^. While the fibroblast network model predicts a number of drugs that modulate fibroblast activation, substantial experimental characterization is needed to capture phenotypic responses to drugs that were not captured by prior modeling.

In this study, we combined high content microscopy, network modeling, and machine learning to identify drugs that differentially regulate fibroblast phenotypic metrics and predict their underlying network mechanisms. We used image-based feature extraction to more deeply characterize drug response and fibroblast phenotype, capturing drug-induced changes across a set of single-cell metrics relevant to fibrosis. Using a novel logic-based mechanistic machine learning approach, LogiMML, we predicted signaling pathways that determine how drugs regulate fibroblast phenotype. Finally, we experimentally validated the main pathway mechanism predicted by the LogiMML model that mediates how Src inhibition suppresses fibrotic responses.

## Results

### An *in vitro* screen for candidate fibrosis drugs

Previously, we applied our published cardiac fibroblast network model^12^ to identify candidate therapies predicted to reduce cardiac fibrosis^13^. This logic-based differential equation network model was developed from a wide range of fibroblast signaling relationships from in vitro studies in the literature. The model predicts changes in fibrotic outputs including collagen I and III, αSMA, EDA fibronectin, matrix metalloproteases, and F-actin in response to changes in extracellular signaling contexts and drug treatment^12^. This model was previously integrated with the drug-target database DrugBank to make predict the response of fibroblasts to 121 FDA-approved or investigational drugs that have targets in this network^13^.

To expand upon the in silico modeling work done in this previous study^13^, we aimed to develop a list of drug candidates to test experimentally for their ability to reduce fibrosis in cardiac fibroblasts in vitro. As the model predicted many drugs to reduce fibrosis to similar quantitative degrees^13^, we included drug selection criteria outside of our modeling results alone to further narrow-down a list of candidate drugs. First, we prioritized pathway diversity of the drug targets to ensure that we would perturb fibrotic signaling comprehensively and avoid testing redundant drugs in our experiments. As drug repurposing has become an increasingly effective and efficient strategy for treating cardiovascular disease, we next looked to prioritize drugs that had previous clinical indications for other disease areas^14, 15^. Using these selection criteria, we developed the following list of thirteen drugs to evaluate experimentally: anakinra, valsartan, defactinib, HW-4-023, glutathione, CW-HM12, salbutamol, marimistat, fasudil, SB203580, pirfenidone, brain natriuretic peptide (BNP), and a combination of valsartan and BNP (Table S1). Among the list of candidate drug targets are regulators for inflammatory signaling, mechanical stretch response, non-canonical TGFβ signaling, and modification of secreted proteins.

We next aimed to test these candidate drugs for their ability to quantitatively reduce fibrosis as characterized by image-based single-cell profiling of procollagen I, α-smooth muscle actin (αSMA), and F-actin. In injury signaling conditions, such as following myocardial infarction (MI), myocardial cells are exposed to elevated proinflammatory^16–18^. To represent these signaling contexts in an in vitro system, we included IL1β and TGFβ, shown to be elevated following cardiac injury, in our treatment conditions to represent proinflammatory and profibrotic contexts respectively^19–21^. We tested our candidate drugs under four total cytokine contexts (baseline context with no added cytokine, fibrotic context represent by TGFβ, inflammatory context represented by IL1β, and combined context represent by both TGFβ and IL1β)^19–21^. In total, we used 108 treatment conditions consisting of one of the thirteen drugs at a low, medium, or high dose combined with one of the four cytokine contexts. We also included treatments of each cytokine context with no drug to establish a control baseline for cell responses to cytokines. We imaged and quantified single-cell protein expression of three fibrotic markers, procollagen I, α-smooth muscle actin (αSMA), and F-actin using high-content microscopy and a custom CellProfiler software pipeline^22^.

Interestingly, the antifibrotic drugs in our screen induced differential effects on fibrosis. Of the 13 candidate drugs, WH-4-023, fasudil, and defactinib caused the strongest reduction of procollagen I, F-actin, and αSMA expression in a TGFβ signaling context, even at the lowest dose (Figure 1A). Conversely, a second set of drugs including anakinra and glutathione increased fibrotic marker expression in both TGFβ and combined TGFβ/IL1β contexts when applied directly to fibroblasts. In a previous clinical study, anakinra, an IL1 receptor inhibitor, was shown to improve cardiac function and prevented heart failure following acute MI^23^. While anakinra has been shown to reduce infarct scar area in a mouse MI model, it also exhibits other beneficial cardiac effects post-MI including inhibition of post-MI myocyte apoptosis and reduction in systemic inflammat^24, 25^. Based on these previous studies, it is likely that anakinra has a net antifibrotic effect on fibroblasts in the presence of other myocardial cell types even though anakinra treatment increased fibrotic marker expression in this experiment. A third set of drugs showed more selective antifibrotic effects. For example, while fasudil significantly reduced expression of all three fibrosis markers in a TGFβ signaling context, pirfenidone only significantly reduced F-actin (Figure 1 B-E). This third set of drugs is of particular interest as it contains drugs that differentially regulate markers for fibrosis. Given the recent clinical effectiveness of pirfenidone for lung fibrosis, and success in diseases models for cardiac fibrosis^26, 27^, we further investigated the mechanisms by which it regulates F-actin in cardiac fibroblasts.

**Figure 1:**
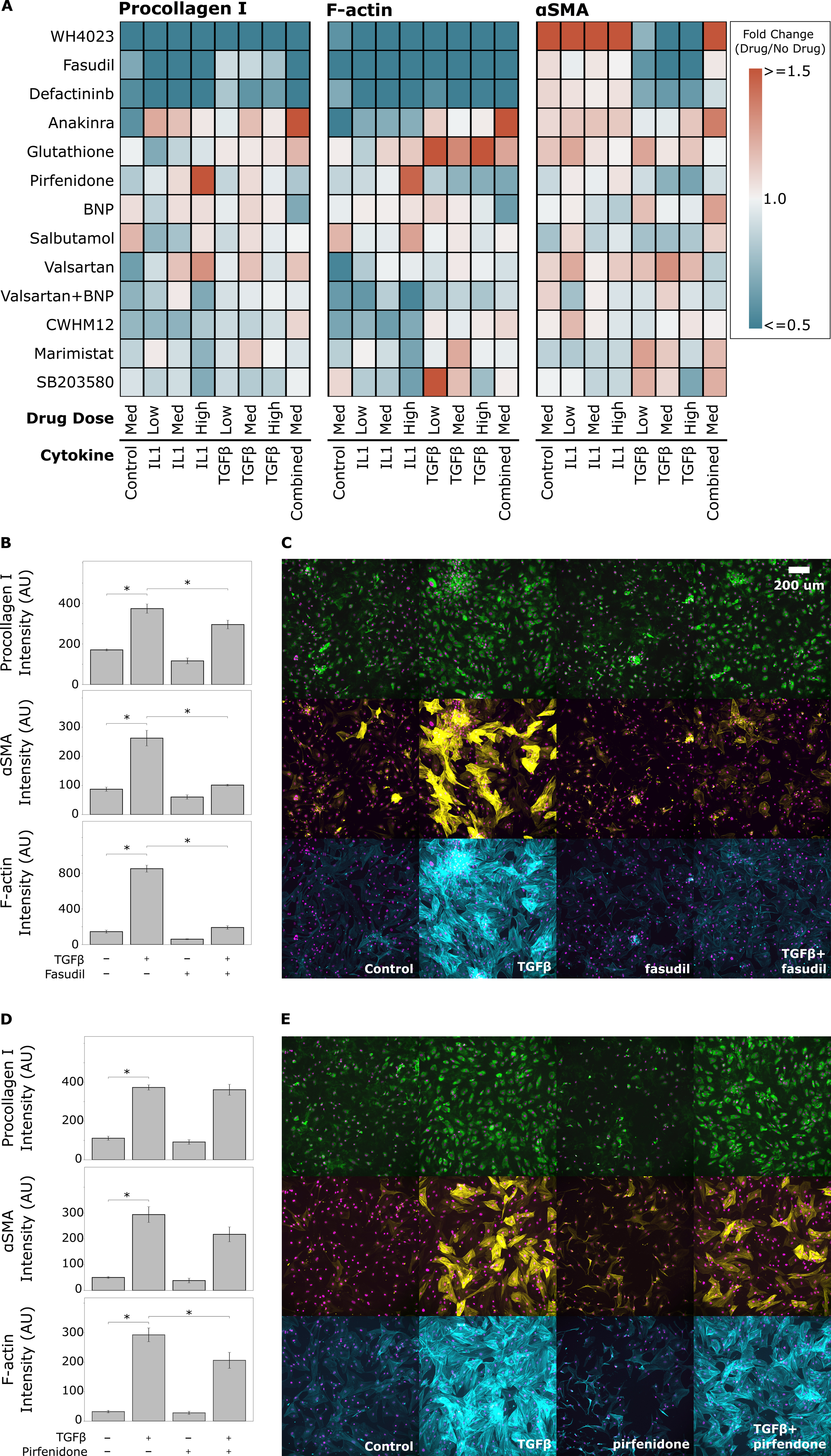
High-content microscopy screen for drugs that module fibroblast activation. A) Expression of fibroblast activation markers procollagen I, F-actin, and αSMA in human cardiac fibroblasts upon treatment of 13 drugs at 3 doses, under environmental contexts of TGFβ, IL1β, or both. Fold change values show ‘drug vs. no drug’ Integrated Intensities for each protein. Panels B and C show quantification and representative images of the effects of 50 µM fasudil, a Rho-kinase inhibitor, which differentially regulates fibrotic protein expression. Panels D and E show quantification and representative images of the effects of 10mg/mL pirfenidone, a non-specific inhibitor of TGFβ expression, which consistently regulates fibrotic protein expression. *p≤0.05 ANOVA with Tukey’s post-hoc.

### LogiMML: logic-based mechanistic machine learning model predicts how drugs regulate fibroblast phenotype

Assembled actin filaments play a key role in contractility as fibroblasts transition to become myofibroblasts^28^. Therefore, we asked whether the previous mechanistic computational model of the fibroblast signaling network^12^ could predict our experimentally measured inhibition of filament assembly by pirfenidone from Figure 1D. While the model had correctly predicted responses to a number of drugs including galunisertib^13^, here, the original mechanistic model did not capture the ability of pirfenidone to suppress actin filament assembly in a TGFβ signaling context (Figure 2 A).

**Figure 2:**
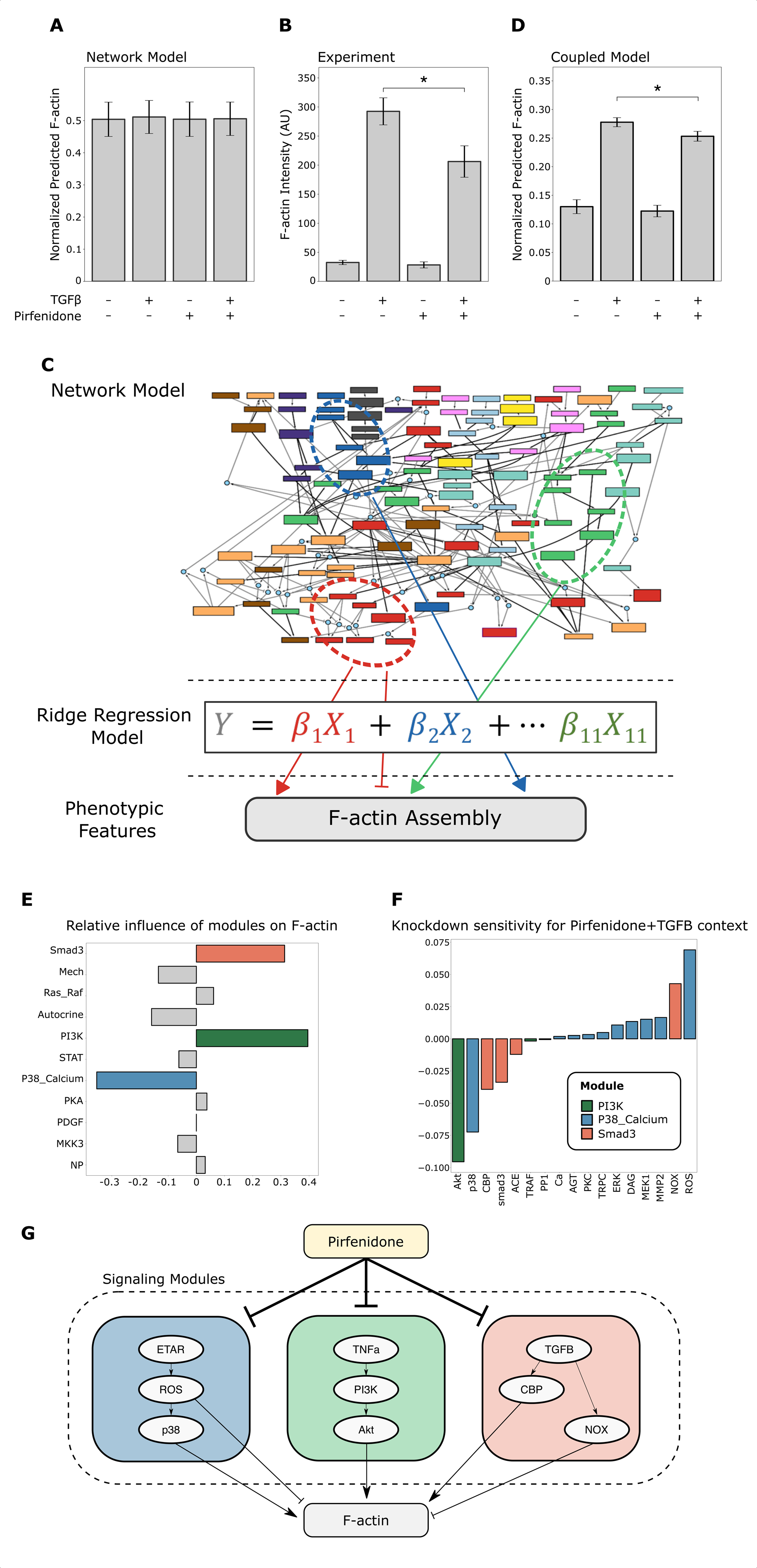
LogiMML logic-based mechanistic machine learning approach guides model revision and predicts network mechanisms underlying pirfenidone suppression of F-actin. A) Original fibroblast network model predicts no change in F-actin upon TGFβ or pirfenidone treatment. Experimental data shows pirfenidone significantly reverses the increase of F-actin by TGFβ (data previously shown in figure 1 D). B) Schematic of the LogiMML approach for integrating logic-based network modeling with machine learning to predict network mechanisms for cell phenotypes. The average activity within each network module is mapped to predict fibroblast phenotypic features via a Ridge regression layer. C) The Coupled LogiMML model predicts TGFβ and pirfenidone effects on F-actin that qualitatively match experimental data shown in panel A. D) LogiMML ridge regression coefficients show predicted relative influence of network modules on F-actin. E) LogiMML node knockdown sensitivity analysis in the context of TGFβ+pirfenidone. Nodes from most influential modules are sequentially knocked down, predicting change in F-actin upon knockdown. F) Schematic of the network mechanisms predicted for the actions of pirfenidone on F-actin, derived from sensitivity analysis in panel E.

Given the limitations of a model based only on prior knowledge, we asked whether drug predictions could be improved by combining the mechanistic model with a machine learning model that leverages data from the drug screen. Motivated by ‘white-box’ machine learning strategies that combine mechanistic models with machine learning^29, 30^, we designed a logic-based mechanistic machine learning (LogiMML) model to predict key regulators that conduct signaling from network model inputs and simulated drugs to experimentally measured phenotypic outputs (Figure 2 B, Figure S 1). As the 108 treatments were insufficient to infer new links to phenotypic outputs from all 91 model nodes, we reduced the model’s dimensionality by clustering nodes into modules. Eleven signaling modules were computed based on a combined influence and sensitivity analysis, grouping nodes with similar predicted behavior across signaling contexts. The machine learning component was then trained by mapping the model-predicted activity of each network module for each of the 108 drug+cytokine treatments to respective experimentally measured outputs. Regularized ridge regression was selected for the machine learning layer of the LogiMML model to reduce the likelihood of overfitting^31^. As measured experimentally, the LogiMML model correctly predicted the respective induction and suppression of F-actin by TGFβ and pirfenidone (Figure 2 C). Leave-one-out cross validation (LOOCV) was performed on the LogiMML model to evaluate performance across variations in the experimental data set. The means and standard deviation of the LOOCV MSE values were 0.022 and 0.080 for the F-actin Integrated Intensity model.

We next asked whether the LogiMML model could provide new mechanistic insights into how F-actin is regulated by pirfenidone. First, we used the LogiMML model’s ridge regression coefficients to predict the modules that most influence F-actin. We used the β coefficients from the LogiMML model to predict the influence of a given signaling module on the cell feature of interest. ‘PI3K’ and ‘Smad3’ modules were predicted to be the top positive regulators of F-actin, while the ‘P38_Calcium’ module was predicted as the top negative regulator (Figure 2 D). These predictions for fibroblasts are consistent with previous studies with other cell types showing that members of the ‘Smad3’ and ‘P38_Calcium’ signaling modules regulate F-actin filament assembly in endothelial cells and that members of the ‘PI3K’ signaling module promote actin filament remodeling during migration in embryonic fibroblasts ^32–34^. To identify which individual signaling nodes within these three modules most regulate F-actin, we performed a virtual knockdown screen of the mechanistic network model for regulators of F-actin in a ‘TGFβ+pirfenidone’ signaling context (Figure 2 E). In these analyses, the quantitative outputs of the model are normalized outputs that can be compared to determine predicted increases or decreases in a cell feature in response to a perturbation. Summarizing these analyses, the LogiMML model predicts that pirfenidone regulation of F-actin is positively regulated by p38, Akt, and CBP, while negatively regulated by ROS and NOX (Figure 2 F).

### Drugs and pathways controlling fibroblast morphology and texture

Given the differential regulation of fibrosis marker protein expression, we asked whether other aspects of fibroblast phenotype may also be differentially regulated by drugs and cytokines. Qualitatively, we observed morphological changes in cell shape, actin-myosin stress fiber formation, intracellular protein distribution, and cell area (e.g. for pirfenidone treatment see Figure 1 E). To measure these characteristics of fibroblast phenotype, we developed a custom CellProfiler image analysis pipeline quantifying 137 total single-cell cell features^22, 35^. Integrated intensities for the three fibrotic marker proteins, procollagen I, F-actin, and αSMA clustered relatively close to each other across the feature space (Figure 3 A). As expected, expression of these marker proteins and similar features were high under TGFβ and TGFβ-like treatments, and low under negative control and IL1β conditions. While the central rows of the heatmap contain many features with similar treatment responses, the features at the top and bottom regions of the heatmap show high heterogeneity in response to drugs. The significance of the overall correlation between actin, aSMA, and collagen expression is two-fold: that some drugs such as fasudil suppress a well-studied canonical myofibroblast activation program, and that the responses to other drugs revealed that the overall phenotype space of fibroblasts is much more diverse and can be specifically targeted with drugs like WH-4-023. Even within the actin/aSMA/collagen cluster, the hierarchical clustering shows some examples where only in the context of TGFβ treatment, some drugs up-regulated aSMA/collagen features while down-regulating actin features.

**Figure 3:**
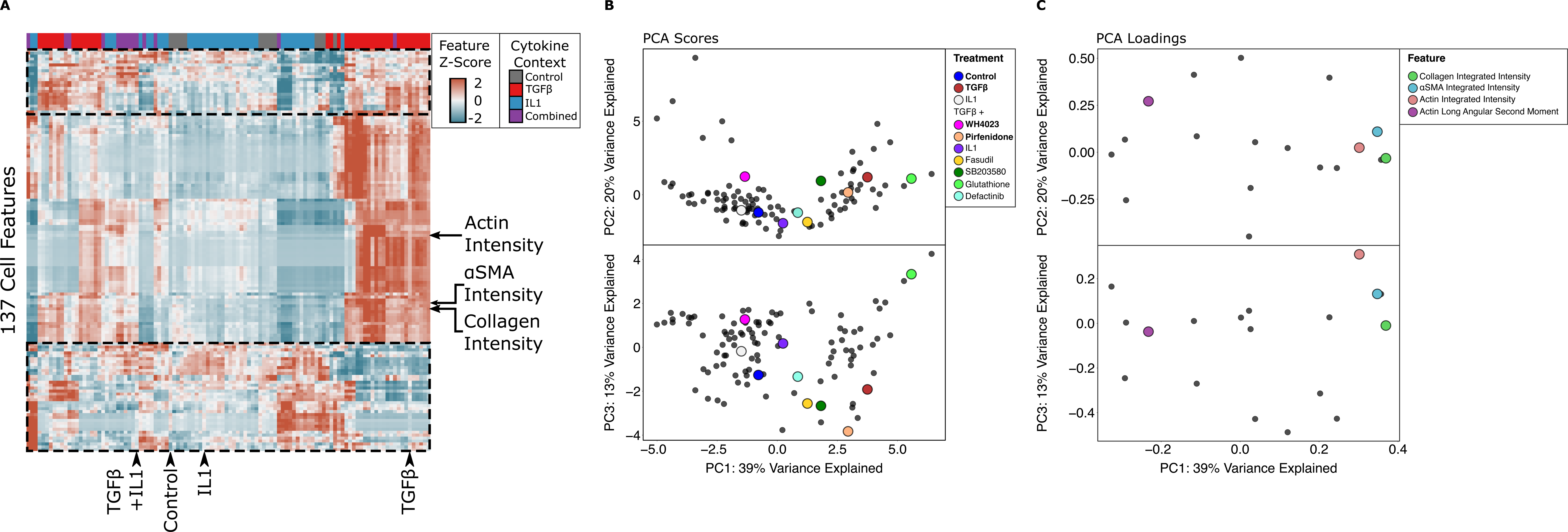
Survey of single-cell fibroblast phenotypic features in response to 13 drugs at 3 doses and 4 environmental contexts. A) 137 single-cell fibroblast features that quantify protein intensity, protein localization, cell morphology, and fiber texture. This heatmap was organized on treatment and feature axes by agglomerative hierarchical clustering. B) Principal component scores of experimental data reduced to a set of 18 representative fibroblast features. C) Principal component loadings the reduced of PCA scores and loadings define a primary axis of fibroblast activation with correlated protein expression of procollagen, αSMA, and F-actin that is modulated by many drugs. Off-axis, the Src inhibitor WH-4-023 modulated the cell texture feature Actin Long Angular Second Moment, which motivated further study.

The extraction of fibrotic marker proteins and the large degree of information about those fibrotic features is a rich dataset with which we next wished to understand more directly how they relate to each other and to treatment. Given the risk that some of our features carry redundant information, we calculated the correlation between all features and clustered the correlation matrix. This identified 15 strong sets of feature clusters. We selected one representative feature of each cluster (Figure S2, Figure S3) based on choosing the cluster member that demonstrated high variance across samples and low correlation with features from other clusters. In addition to the 15 representative features, we retained the three integrated intensity features for procollagen I, F-actin, and αSMA, yielding a final set of 18 distinct features of the original 137 features, which have the potential to represent the complexity of the larger dataset.

In order to interpret the overall underlying relationships in the 18 selected features and how they relate to treatments, we performed Principal Component Analysis (PCA) (Figure 3B-C). Negative control treatments had a negative score on the first principal component (PC1), while cells treated with TGFβ showed a high positive score on PC1, indicating that the first principal component correlates with an axis of classical fibroblast activation (Figure 4 B, Figure S4 A). This was further supported by the PCA loading values for integrated procollagen I, F-actin, and αSMA (Figure 3 C, Figure S4 B). These three features are expected to be relatively high in activated myofibroblasts and indeed have strong positive loadings on PC1. On the PCA scores, many of the ‘TGFβ + Drug’ groups deviated from the control-TGFβ axis defined on PC1, implying that drugs induce phenotypic changes distinct from a simple reversal of TGFβ’s effects. To further investigate drug-induced changes in phenotype, we analyzed the PCA scores and loadings to infer links between drugs and the features they regulate. Notably, the Src inhibitor WH-4-023 (WH) showed directionality on the scores plot similar to that of Actin Long Angular Second Moment (Actin Long ASM, a measure for actin uniformity) on the loadings plot. Actin-myosin stress fibers, composed of multiple actin filaments along with other proteins, contribute to pathological fibrosis and myofibroblast differentiation^36–38^. This feature and treatment pair showed a negative value on PC1 and a positive value on PC2 relative to the TGFβ and control groups, respectively. The similar directionality of WH and Actin Long ASM suggests that Src inhibition may modulate actin uniformity.

**Figure 4:**
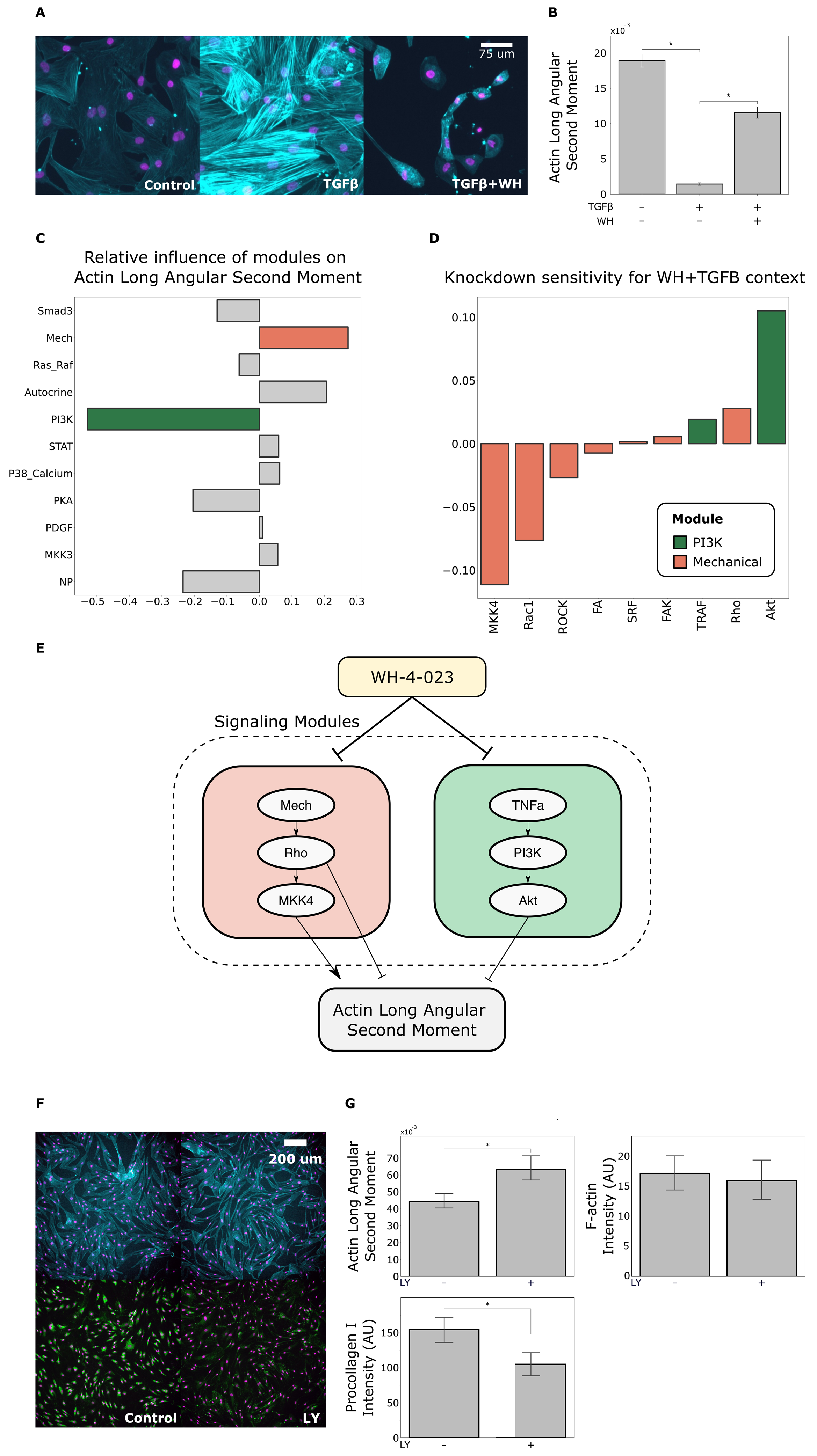
Logic-based mechanistic machine learning predicts the PI3K module to mediat how Src inhibitor suppresses stress fibers, validated by subsequent experiments. A) Images of human cardiac fibroblasts treated with baseline control stimulus, TGFβ, or TGFβ + 20 µM WH-4-023. B) Quantification of Actin Long Angular Second Moment (ASM), a measure of actin uniformity and reduced stress fibers based on images in panel A. C) Regression coefficients from the LogiMML mechanistic machine learning model that predicts network modules that regulate actin long ASM. D) Knockdown sensitivity analysis predicting individual proteins that regulate actin long ASM in the TGFβ+WH-4-023 signaling context. E) Signaling schematic for WH-4-023 effect on actin long ASM, derived from sensitivity analysis in panel D. F) Human cardiac fibroblasts treated with 20 µM PI3K inhibitor LY294002 or baseline control stimulus, measuring F-actin and procollagen expression. G) Quantification of long actin Angular Second Moment (measure of actin uniformity), F-actin integrated intensity, and Procollagen I integrated intensity. *p≤0.05 ANOVA with Tukey’s post-hoc in panel B, and *p≤0.05 Student’s T-test in panel G.

Based on the initial inference from the PCA, we revisited the images from the high-content microscopy experiment. Fibroblasts treated with TGFβ exhibited discrete actin-myosin stress fibers, and stress fibers were qualitatively reduced when WH-4-023 (WH) was added (Figure 4 A). Quantitative analysis of actin uniformity (inversely correlated with stress fibers) using Actin Long Angular Second Moment (ASM) further supported that TGFβ increased and Src inhibitor WH reduced actin uniformity (Figure 4 B). The full dose response for Long Actin ASM to WH-4-023 is shown in Figure S5.

To predict the signaling pathways that specifically regulate actin-myosin stress fibers, we again applied the LogiMML coupled modeling approach, but this time training the ridge regression layer of the model on experimental measurements of Actin Long ASM. The means and standard deviation of the LOOCV MSE values were 0.083 and 0.142 for the Long Actin ASM model. The LogiMML model regression coefficients predicted that the ‘Mechanical’ module was the top positive regulator of Actin Long ASM and that the ‘PI3K’ module was the top negative regulator of Actin Long ASM (Figure 4 C). To identify which individual signaling nodes within these two modules most regulate Actin Long ASM, we performed a virtual knockdown screen of the mechanistic network model for regulators of Actin Long ASM in the context of ‘TGFβ+WH-4-023’ and predicted that Rho, MKK4, and Akt are proximal regulators of Actin Long ASM and actin-myosin stress fiber formation (Figure 4 D-E).

### PI3K signaling stimulates actin-myosin stress fiber formation and collagen expression

After deriving a putative signaling schematic for Actin Long ASM using the LogiMML model, we aimed to experimentally validate the prediction that inhibition of PI3K/Akt would suppress stress fiber formation and thereby increase Actin Long ASM (Figure 4 E). In previous studies using PI3K inhibitors, PI3K was shown to regulate fibroblast contractility, fibroblast-to-myofibroblast transition, and TGFβ-induced αSMA and collagen production^39, 40^. Given these previously implicated roles for PI3K in myofibroblast activation and fibrosis, we wanted to investigate if PI3K has a regulatory role for actin-myosin stress fiber formation in cardiac fibroblasts. We treated human cardiac fibroblasts with either a negative control condition or a 20 µM dose of the PI3K inhibitor LY294002 (LY). Treatment with LY significantly increased Actin Long ASM, but notably, it had no significant effect on the total assembly of actin filaments in each cell, measured by integrated F-actin intensity (Figure 4 F-G). This selective effect of PI3K inhibition on stress fiber formation, while having no significant effect on total F-actin, suggests that actin filament assembly and stress fiber formation are differentially regulated processes. PI3K inhibition also significantly reduced integrated procollagen I intensity, demonstrating a role for PI3K signaling in cardiac fibroblast collagen production (Figure 4G).

## Discussion

Cardiac fibroblasts are central regulators and promising therapeutic targets following cardiac injury. To identify how clinically relevant drugs regulate diverse aspects of fibroblast phenotype, we performed high-content screening of 13 drugs in 4 environmental contexts. We expanded our high-content microscopy feature set to 137 single-cell features, measuring fibrotic marker protein intensity, intracellular protein distribution, fiber texture, and cell morphology. After reducing the feature space and dimensionality of our experimental data, we found that many aspects of fibroblast phenotype are uniquely induced by drug and cytokine treatments. Notably, when administered with TGFβ, the drugs WH-4-023, defactinib, fasudil, and pirfenidone induced phenotypes that deviated from the PCA axis corresponding to classical TGFβ response. The differences between these phenotypes can be partially explained by differential drug regulation of features capturing Procollagen I and αSMA expression, and actin filament assembly and actin-myosin stress fiber formation. To predict how drugs regulate cell signaling and influence phenotype, we developed the logic-based mechanistic machine learning (LogiMML) approach which coupled the logic-based fibroblast network model with a ridge regression model trained on the high-content drug screen. Using this expanded LogiMML model, we predicted regulatory mechanisms for pirfenidone and Src inhibitor WH-4-023 on actin filaments. We predicted that pirfenidone regulates actin filament assembly via the ‘P38_Calcium’, ‘Smad3’, and ‘PI3K’ signaling modules, with Akt, p38, and CBP predicted to be positive drivers of actin filament assembly within these modules. We also predicted that WH-4-023 regulates actin-myosin stress fiber formation via the ‘PI3k’ and ‘Mechanical’ signaling modules. As predicted by the LogiMML model, we experimentally validated that PI3K inhibition reduces actin-myosin stress fiber formation in human cardiac fibroblasts. These studies validate the ability of the LogiMML approach to predict signaling mechanisms from a phenotypic screen.

### Differential regulation of fibroblast phenotype by drugs and the development of targeted antifibrotic therapies

Drugs that specifically target fibroblast signaling may provide directed control over the fibrotic response. A major challenge in therapeutic development for fibrosis is that many drugs capable of reducing fibrosis target non-specific regulatory pathways outside of the fibrotic response. For example, the ALK5 inhibitor galunisertib targets the TGFβ receptor and shows promising therapeutic reduction of fibrosis across organs^41–43^. While TGFβ receptor inhibition can reduce fibrosis, recent efforts in target discovery have successfully identified new approaches to mitigate fibrosis that are more fibroblast specific. For example, it was shown that activating fibroblast-specific TLR4 in mice can drive the development of skin and lung fibrosis and that TLR4 inhibition reduces αSMA expression and collagen production in fibroblasts ^44^. Another study showed that fibroblast-specific knockout of STAT3 ameliorates skin fibrosis, and that pharmacological inhibition of STAT3 successfully reduces myofibroblast activation, collagen accumulation, and dermal thickening in experimental fibrosis in mice^45^. Future work can advance our understanding of how candidate drugs regulate specific components of the fibrotic response in fibroblasts and provide targeted control of fibrosis.

### Features of cardiac fibroblast phenotype

Traditional fibrotic markers are not always expressed in fibroblasts and exhibit significant heterogeneity and context dependence, in this study we aimed to explore multiple features of the fibrotic response^46^. Following the reduction of the original set of 137 single-cell features from our high content image analysis, we identified a set of 18 phenotypic features of fibroblasts that exhibit high heterogeneity in response to drug treatments (Figure S 3, Table S 2). Notably, many of the features measuring fiber texture for αSMA and F-actin show different response patterns compared to features measuring overall expression level for those respective proteins (i.e. aSMA integrated intensity versus aSMA long correlation). This distribution of features indicates that the expression and organization of aSMA and actin filaments are independently regulated by candidate drugs. The processes of αSMA protein expression and fiber assembly have different degrees of contribution to pathological fibrosis. For example, a recent study showed that fibroblasts can compensate for the loss of Acta2 transcription and form stress fibers using similar proteins, implying that stress fiber formation is more important than αSMA production for the fibrotic response^47^. Incorporating an expanded set of measurements in future fibrosis studies may provide greater resolution of the fibrotic phenotype in response to therapies and help evaluate changes in pathologically relevant features beyond protein expression.

### Contributions of the LogiMML mechanistic machine learning approach

Mechanistic logic-based differential equation models have enabled systematic prediction of drug action, yet these models are limited by the availability of priori knowledge ^13, 48–50^. An alternative is machine learning, although ‘black-box’ ML approaches like artificial neural networks predict input-output relationships without mechanistic insight. In contrast, two recent studies combined mechanistic modeling with machine learning models like regression and visible neural networks to predict antibiotic stress on metabolism and drug synergies for cancer^29, 51^. These ‘white-box’ approaches provide greater transparency of the intermediate layers between input and output^52^.

The prior approach most similar to the LogiMML framework is ‘white-box’ machine learning proposed by Yang et. al^53^. In that study, a flux balance model of E. Coli metabolism with simple linear regression to predict metabolic reactions important to growth on particular carbon sources. In this study, we propose a method that combines regularized regression with a logic-based model to predict signaling pathways in response to signaling perturbations representative of drug effects.

Building on such advances for logic-based biological networks, our LogiMML mechanistic machine learning approach combines the flexible trainability of a machine learning model with the robust experimentally-determined internal network structure of a mechanistic model. In this study, we used the LogiMML model to predict signaling mechanisms that mediate how drugs regulate F-actin assembly and stress fiber formation in cardiac fibroblasts. However, this is just one of many possible applications for this modeling framework. The LogiMML approach is designed to work across multiple mechanistic modeling formalisms and types of experimental data, coupling the mechanistic model and data to predict mechanisms for the phenotype of interest. The flexible nature of LogiMML presents promising future applications to elucidate cell signaling across many disease areas.

### Src kinase as a therapeutic target for fibrosis

Of the 13 drugs used in this study, the Src inhibitor WH-4-023 (WH) was one of three drugs that showed a strong reversal in TGFβ-induced actin filament assembly, αSMA, and procollagen I expression. WH was also effective at reversing the formation of actin-myosin stress fibers in response to TGFβ. Src inhibitors dasatinib, ponatinib, and saracatinib have all been used in clinical trials across different types of cancer^54–58^. In cancer, Src has been shown to promote proliferation and metastasis through many signaling targets including FAK, Akt, Ras, and PI3K^59–63^

Given that Src signaling affects many central regulatory pathways, recent studies have tested the potential for Src inhibition as a therapy for fibrotic disease. In a renal fibrosis study, blocking Src kinase using PP1 was shown to inhibit TGFβ-induced expression of collagen I, αSMA, and fibronectin^64^. In that study, Src inhibition was also shown the reduce the development of renal fibrosis in obstructed kidneys in vivo in mice, indicating Src inhibition as a potential renal fibrosis and chronic kidney disease therapy. Another study focusing on lung fibrosis showed that TGFβ induces Src kinase activity in lung fibroblasts and that Src is required for myofibroblast contraction^65^. Further, inhibition of Src kinase in vivo with AZD0530 reduced scar area and αSMA expression in mice with bleomycin-induced lung fibrosis^65^.

While PI3K signaling has established relevance in fibrotic pathologies, this signaling pathway has been heavily implicated in other disease models including cancer. Aberrant activation of PI3K signaling has been shown to contribute to tumor progression in multiple cancers including breast, lung, and ovarian cancers^66^. Changes in extracellular matrix remodeling have been shown to influence many classically defined hallmarks of cancer, with ECM adhesion-induced PI3K signaling shown to regulate self-sufficient cell growth via FAK signaling^67, 68^ . Given the large overlap between regulators of cytoskeletal restructuring and tumor progression, therapeutic targets like PI3K could be efficient future drug targets that can modulate pathways governing multiple diseases. The interplay between cytoskeleton regulation and cancer progression should be further explored to identify other central regulators that may modulate both cancer and fibrotic disease progression.

In this study, we applied the LogiMML network to investigate how Src contributes to actin-myosin stress fiber formation induced by TGFβ. We predicted that PI3K signaling contributes to profibrotic Src signaling in cardiac fibrosis. This proposed mechanism is supported by previous studies, showing that PI3K regulates fibroblast contractility and myofibroblast activation in skin fibroblasts, and TGFβ-induced αSMA and collagen production in lung fibroblasts^39, 40^. To validate this proposed profibrotic role for PI3K, we show that PI3K inhibition reduced procollagen I production and actin-myosin stress fiber organization in HCFs. While previous work has shown that mechanical stretch, Rho-kinase, and myosin light chain kinase (MLCK) positively regulate the organization of actin filaments into stress fibers, the role of PI3K’s regulation of actin-myosin stress fiber formation has not been thoroughly explored^69, 70^. Here, we show that treatment with PI3K inhibitor LY294002 (LY) significantly reduces stress fiber formation without affecting the total amount of assembled actin filaments, implying PI3K positively and specifically regulates actin-myosin stress fiber formation in cardiac fibroblasts. Future studies should explore if Src kinase inhibitors mitigate cardiac fibrosis in vivo, and to what degree PI3K kinase contributes to the regulation of cardiac fibrosis by Src.

### Limitations and future directions

The main limitation of this study is that our modeling and experimental approaches address cell signaling in cardiac fibroblasts in vitro, but do not address how fibroblasts respond to drugs in an in vivo signaling environment. Our experimental data also captures some key fibrotic proteins, but does not measure other fibrotic outputs of interest, like EDA fibronectin, and does not capture a comprehensive signaling profile of the fibroblast. Despite these limitations, the LogiMML framework was sufficient to predict a validated role for PI3K in promoting stress fiber formation. Experimentally, future work could include proteomics or RNA-seq analysis of fibroblasts to measure how drugs differentially regulate intracellular molecular profiles. To maximize reproducibility across the drug screen, we used 2D culture on multi-well plates treated with CellBind. More focused follow-on studies could perform secondary validations with various extracellular matrix, stiffness or mechanical stretch. Future modeling work could include simulated conditions for in vivo or in vitro co-culture conditions to incorporate the signaling influence of other cell types. Given the flexibility of the LogiMML modeling approach, these simulated data could be feasibly paired with respective experimental data to make predictions for fibroblast signaling under new conditions.

### Conclusions

In this study, we showed that drugs exhibit differential effects on cardiac fibroblast phenotype and work via distinct mechanisms that can be predicted by logic-based mechanistic machine learning. By expanding the microscopy feature set in the high content imaging pipeline, we captured greater resolution of the fibroblast phenotype and measured how phenotypic features changed in response to drugs. Using our LogiMML modeling approach, we predicted signaling mechanisms for how pirfenidone and Src inhibitor WH-4-023 affect actin filament assembly and actin-myosin stress fiber formation, respectively. We predicted that PI3K regulates F-actin stress fiber formation, which we experimentally validated in human cardiac fibroblasts. This study presents new features of fibroblast phenotype to be further explored in fibrosis, identifies specific roles for PI3K in cardiac fibroblast signaling, and demonstrates an adaptable mechanistic machine learning approach to predict signaling outcomes for fibrosis that can be expanded to other diseases.

## Methods

### *In vitro* experiments in human cardiac fibroblasts

Primary human ventricular cardiac fibroblasts were purchased from PromoCell (PromoCell C-12375; PromoCell GmbH, Germany). Cells were cultured in DMEM containing 10% FBS and 1% Pen/Strep, and were kept in an incubator maintained at 5% CO_2_. Cells were plated in a 96-well plate at 5,000 cells/well and then grown in 10% FBS for 24 hours, serum starved for 24 hours, and then treated with the following cytokine conditions for 96 hours: 0% FBS control media, 0% FBS media with 20ng/mL TGFβ1 (Cell Signaling Technology, 8915LC), and 0% FBS media with 10 ng/mL human IL1β (Cell Signaling Technology, 8900SC), or TGFβ1 and IL1β combined. Cells were treated with these conditions either alone or with 1 of 13 compounds at 1 of 3 concentrations. We determined drug concentrations via a literature search, prioritizing concentrations that yielded significant effects in vitro in fibroblasts or similar cell types. The drugs with their respective concentrations are as follows: [0.25,1,2] µg/ml of anakinra (Kineret, SOBI Inc.), [1,5,10] µM valsartan (Sigma-Aldrich, SML0142-10MG), [0.2,1,2] µM BNP (Sigma-Aldrich, B5900-.5MG), [1,5,10]µM valsartan combos respectively with [0.2,1,2] µM BNP, [10,30,60]mM glutathione (Sigma-Aldrich, G4251-1G), [1,3,5] µM CW-HM12 (Cayman Chemical Company, 19480), [10,20,50] µM salbutamol (Sigma-Aldrich, S8260-25MG), [5,10,25] µM marimistat (Sigma-Aldrich, M2699-5Mg), [1,5,10] µM galunisertib (Selleck Chemicals, S2230), [12.5,25,50] µM fasudil (Sigma-Aldrich, CDS021620-10MG), [10,25,50]µM SB203580 (Sigma-Aldrich, S8307-1MG), [1,5,10] mg/mL pirfenidone (Sigma-Aldrich, P2116-10MG), [5,10,20] µM defactinib (MedChem Express, HY-12289A), [5,10,20] µM WH-4-023 (Sigma-Aldrich, SML1334-5MG), and 20 µM LY294002 (Selleck Chemicals, S1105). Cells were grown in these conditions for 72 hours.

Cells were then fixed in 4% PFA in PBS for 30 minutes, permeabilized and blocked for 1 hour in a solution containing 3% BSA and 0.2% Triton, and then stained overnight at 4°C with a 1:500 Anti-Collagen I antibody (Abcam, ab34710). After overnight incubation, cells were washed 3x in PBS and stained with 1:5000 Dapi, 1:1000 Phalloidin CruzFluor 647 Conjugate (Santa Cruz Biotechnology, sc-363797), 1:250 α-Smooth Muscle Actin antibody (Sigma-Aldrich, C6198), and 1:1000 Goat-anti-Rabbit (ThermoFisher Scientific, A-11034).

### Microscopy and single-cell quantification

96-well plates we imaged using the Operetta CLS High-Content Analysis System (Perkin Elmer). All three replicate wells for each condition were imaged and quantified. To quantify αSMA expression, an automated image analysis pipeline was employed in CellProfiler (Broad Institute)^22^. Fibroblast nuclei were identified by the DAPI signal. Next, the collagen-positive region corresponding to each nucleus was segmented using the “propagate” algorithm, using the segmented nucleus as the seed. Next, Fibroblast boundaries were segmented using the “propagate” algorithm, musing the segmented collagen region as the seed. αSMA signal was integrated within each cell’s boundary. Short, medium, and long texture feature information was derived using the MeasureTexture module in CellProflier using texture scales of 2, 6, and 10 pixels respectively. Texture feature values were calculated by subtracting the smallest angle value of a given feature from the largest angle value of that same feature for each cell. F-actin and procollagen expressions were quantified similarly.

### Statistics

Feature values for each well were determined by taking the median value of the feature across all cells in the center tile of each well. Well median values were used as replicates (n=3). Significance was determined using an ANOVA with Tukey’s posthoc in comparisons between more than two groups, and Student’s T-test in comparisons between two groups. Automated data analysis and statistical calculations were performed using Python 3.8.5 and the ‘statsmodels’ Python module version 0.13.2.

### Model Simulations

Drug simulations in the fibroblast network model were performed as previously described using MATLAB version 2022a^12, 13, 71^. Predicted node activity is calculated using logic-based Hill differential equations. Agonist and antagonist drug relationships were represented by altering the activation function of the target node, representing either competitive or non-competitive drug interactions with the respective target. To better represent the cell-to-cell variability observed in in vitro cell responses to treatments, we employed a previously developed ensemble modeling approach combining multiple simulations with random normally distributed parameters^71^. Ensemble simulations were performed using the MATLAB ‘normrnd’ function from the ‘Statistics and Machine Learning’ toolbox to randomly sample parameters within a normal distribution and simulation n of 100. The randomly sampled parameters and means of the sampling ranges are as follows: baseline ligand inputs (0.25), mechanical input (0.85), drug dose (0.85), and raised ligand inputs (0.6). The sampling range for each parameter was calculated by paramMean + CDV * paramMean where COV=0.0396. This COV value, used to scale stochasticity in the model was determined by taking the average coefficient of variation in F-actin, procollagen I, and αSMA expression in human cardiac fibroblasts treated with TGFβ from our in vitro experiments. Code for all modeling, regression and data analysis is available at at https://github.com/andersnelson/Logic-based_MML.

### LogiMML Network-Regression Coupling

The LogiMML mechanistic machine learning model is comprised of a network model layer and a Ridge regression layer. The independent ‘X’ variables used to train the regression model are node activity values from the network model predicted under each simulated drug and environmental condition. To reduce model complexity, network nodes were clustered into 11 signaling modules derived from k-means clustering on a combined sensitivity and influence analysis on the network model^12^. Sensitivity analysis was performed by systematically perturbing individual node values and measuring the change in all other nodes in response to the perturbed node. The influence matrix, the transposition of the sensitivity matrix, was combined with the sensitivity matrix and this combined matrix was used for the k-means clustering. The node activity values were averaged within each module, and these modules’ mean activity values were fed into the regression layer. The dependent ‘Y’ variables for this model were experimentally measured values from our high-content imaging experiments in human cardiac fibroblasts. Sensitivity knockout analysis was performed by simulating a given drug and cytokine context int network model i.e. ‘TGFβ+pirfenidone’ and sequentially setting each node ymax value to 0, measuring reduction or increase in the dependent variable e.g. ‘F-actin Intensity’ upon knockdown.

## Acknowledgements

We acknowledge Dr. Sarah Ewald at the University of Virginia Carter Immunology Center for providing anakinra used in this study.

