## Supplemental Figures and Tables for "Logic-based mechanistic machine learning on high-content images reveals how drugs differentially regulate cardiac fibroblasts"

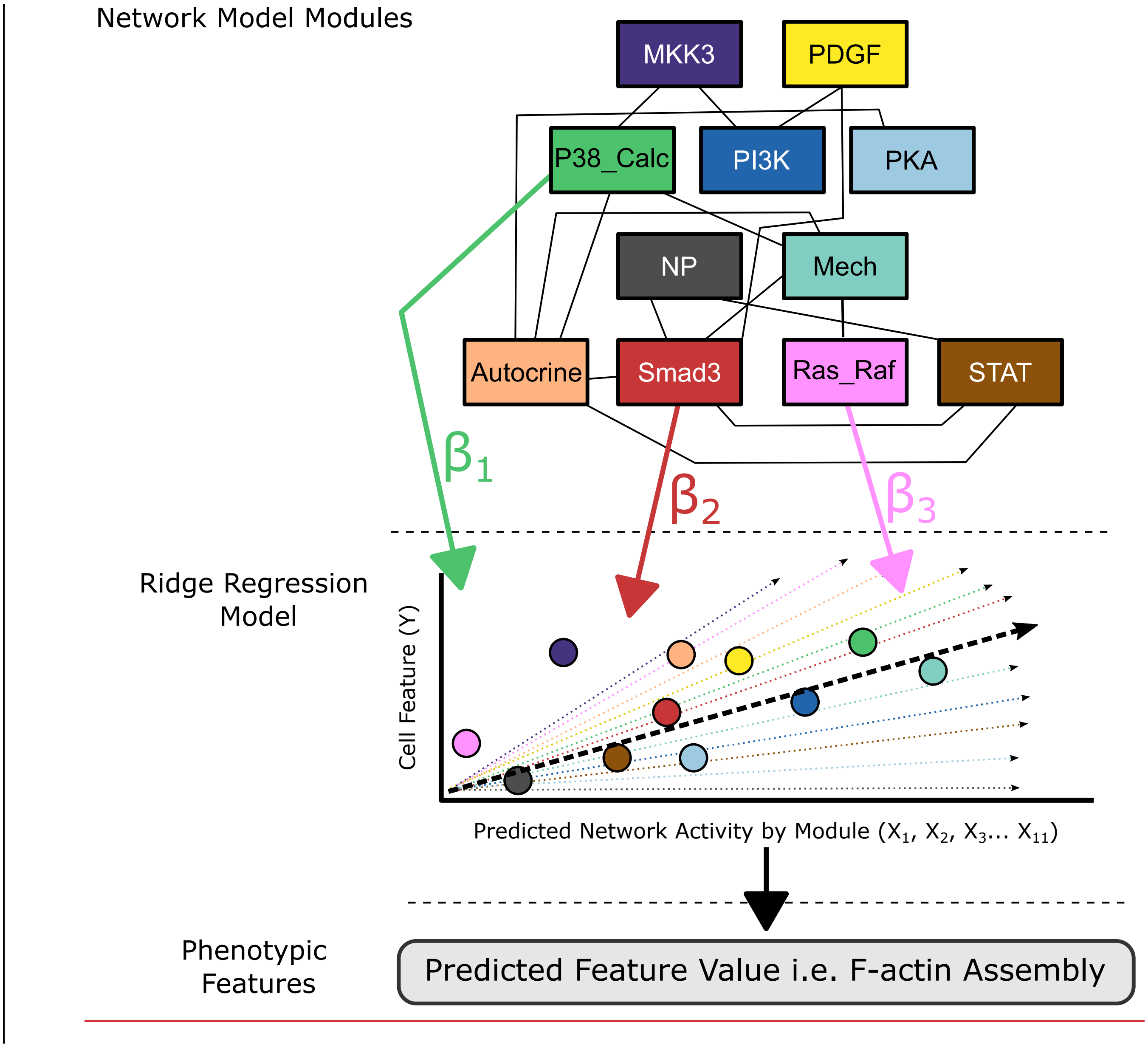


Figure S1: Visual schematic for development of the LogiMML mechanistic machine learning model.

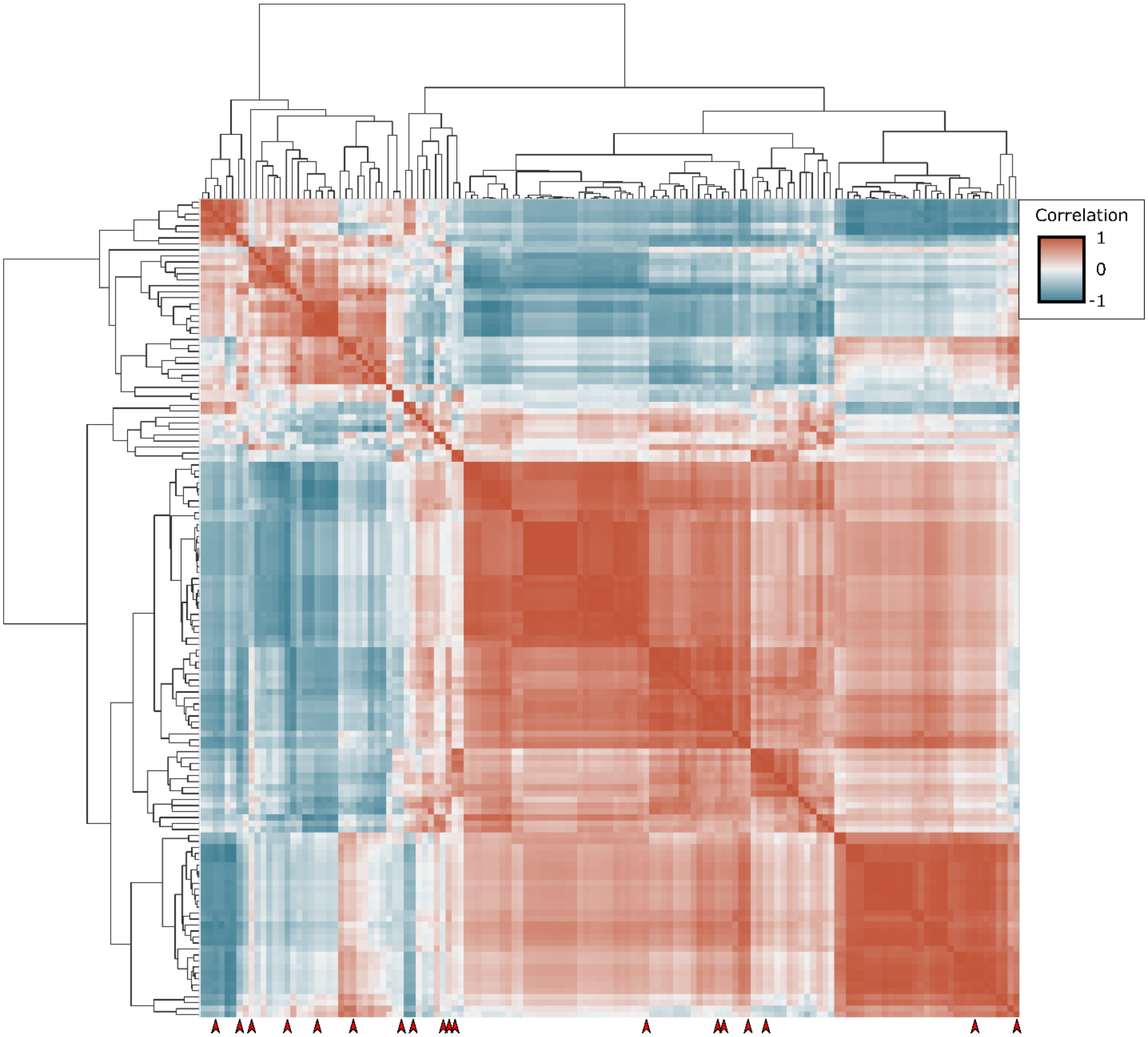


Figure S2: Correlation matrix of the full high-content imaging feature set. The matrix is clustered using agglomerative hierarchical clustering into 15 clusters. Features that were retained in the reduced feature set are denoted by red arrows.

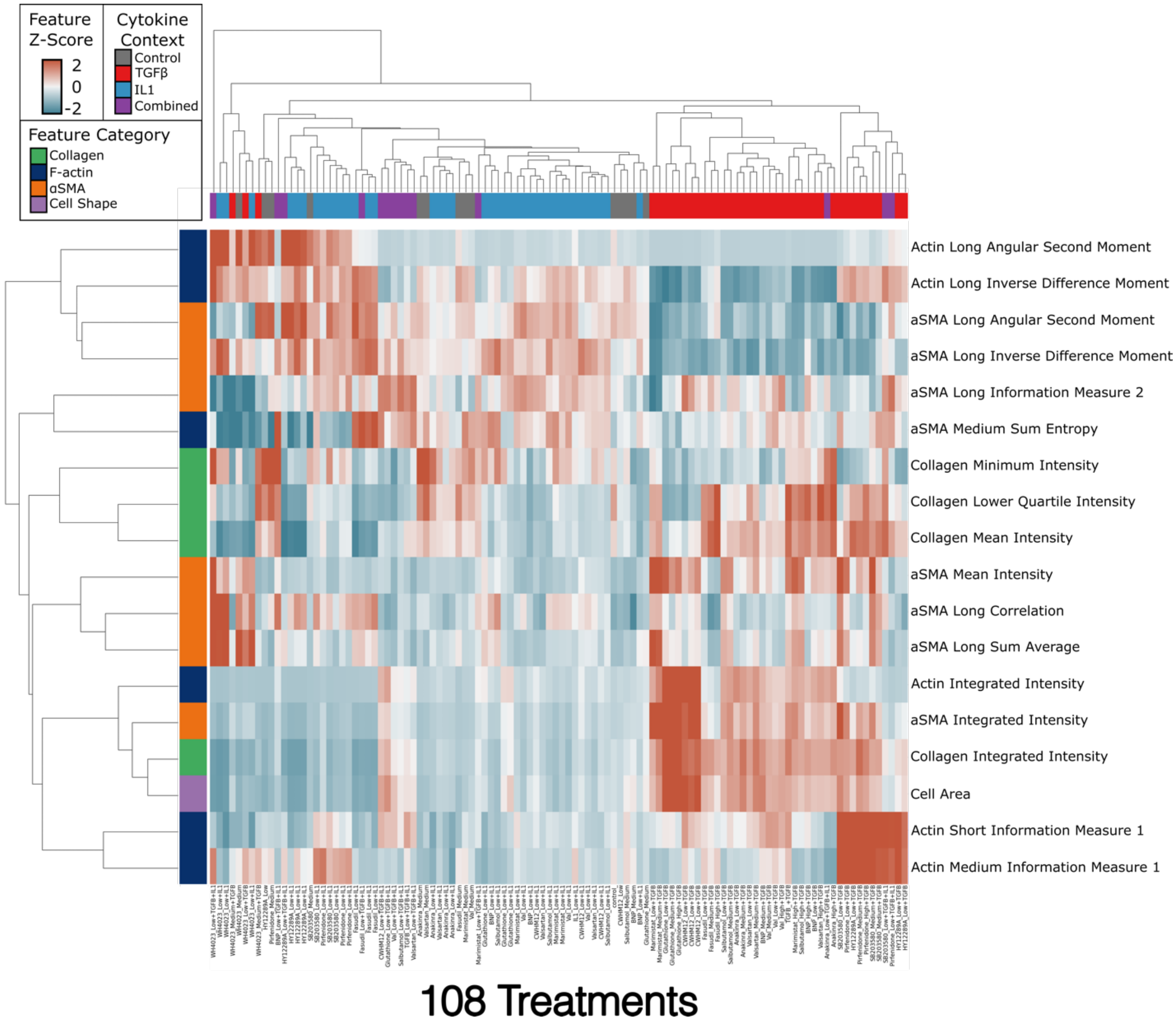


Figure S3: Heatmap of reduced features shown in figure 3C, with all 108 treatment groups shown. Features are clustered by hierarchical agglomerative clustering.

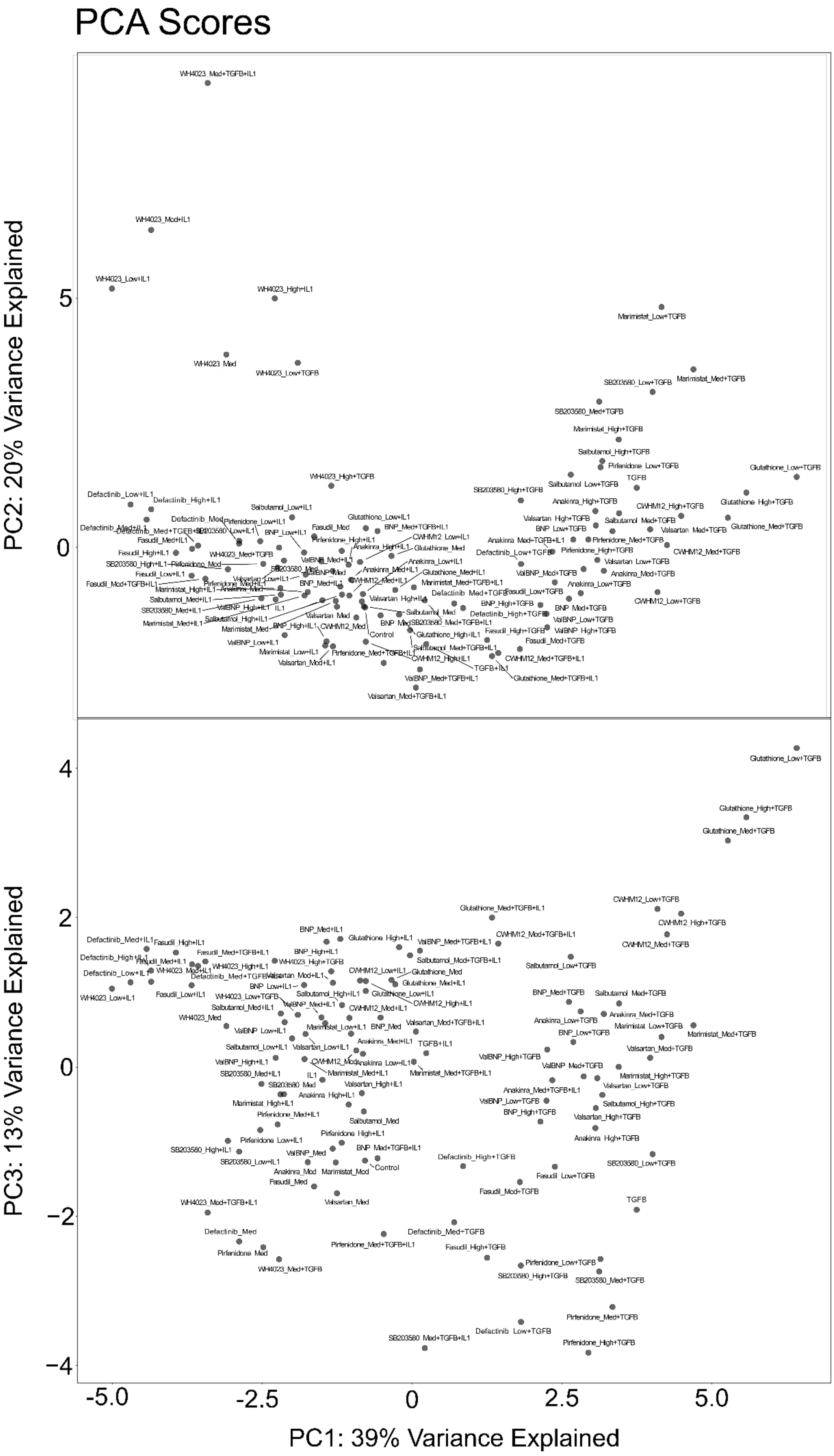
PCA Scores
PC2: 20% Variance Explained
PC3: 13% Variance Explained
PC1: 39% Variance Explained
5
0
4
2
0
−2
−4
-5.0
-2.5
0
2.5
5.0

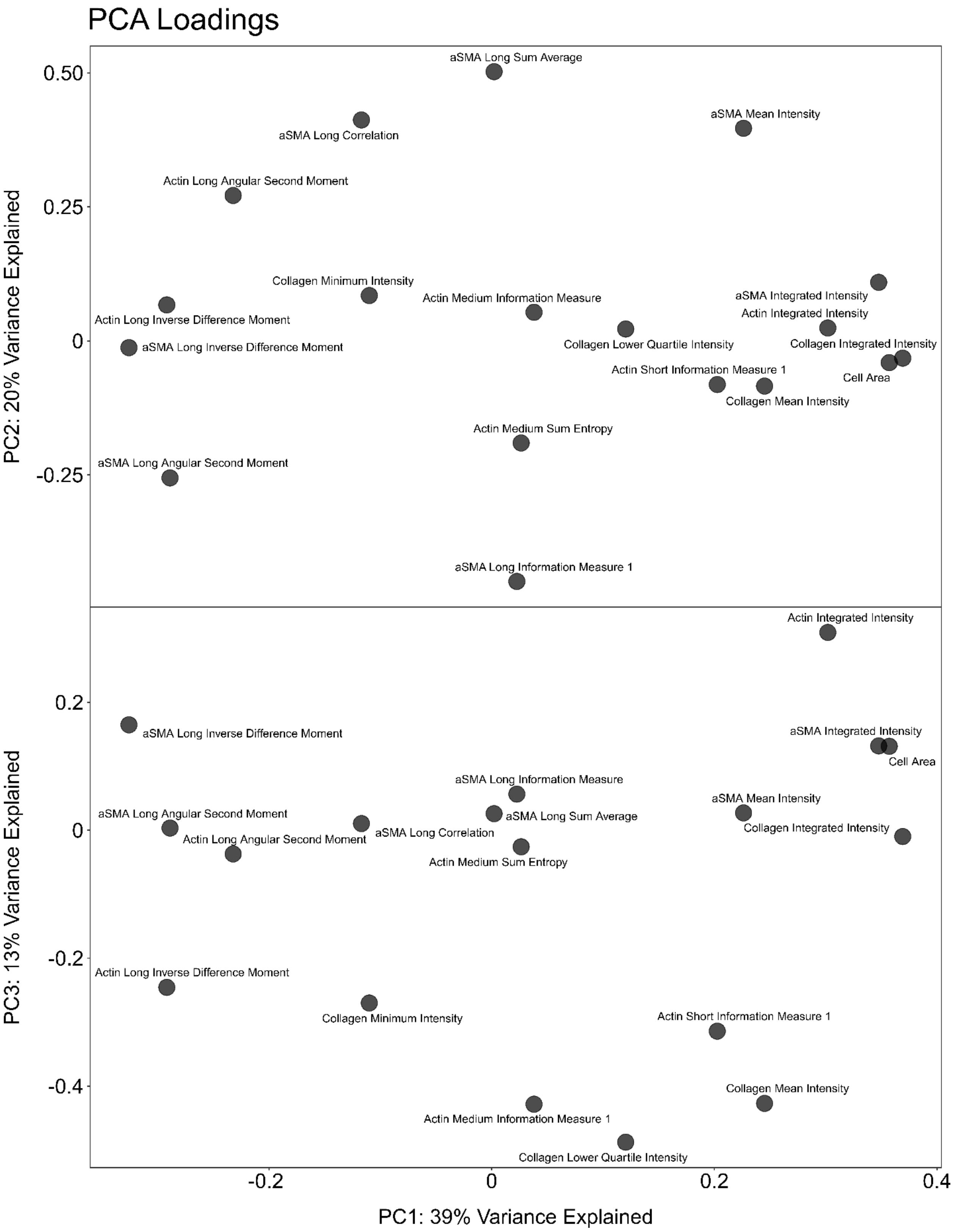


Figure S4: Fully annotated PCA projection of 108 treatments and 18 phenotypic features.

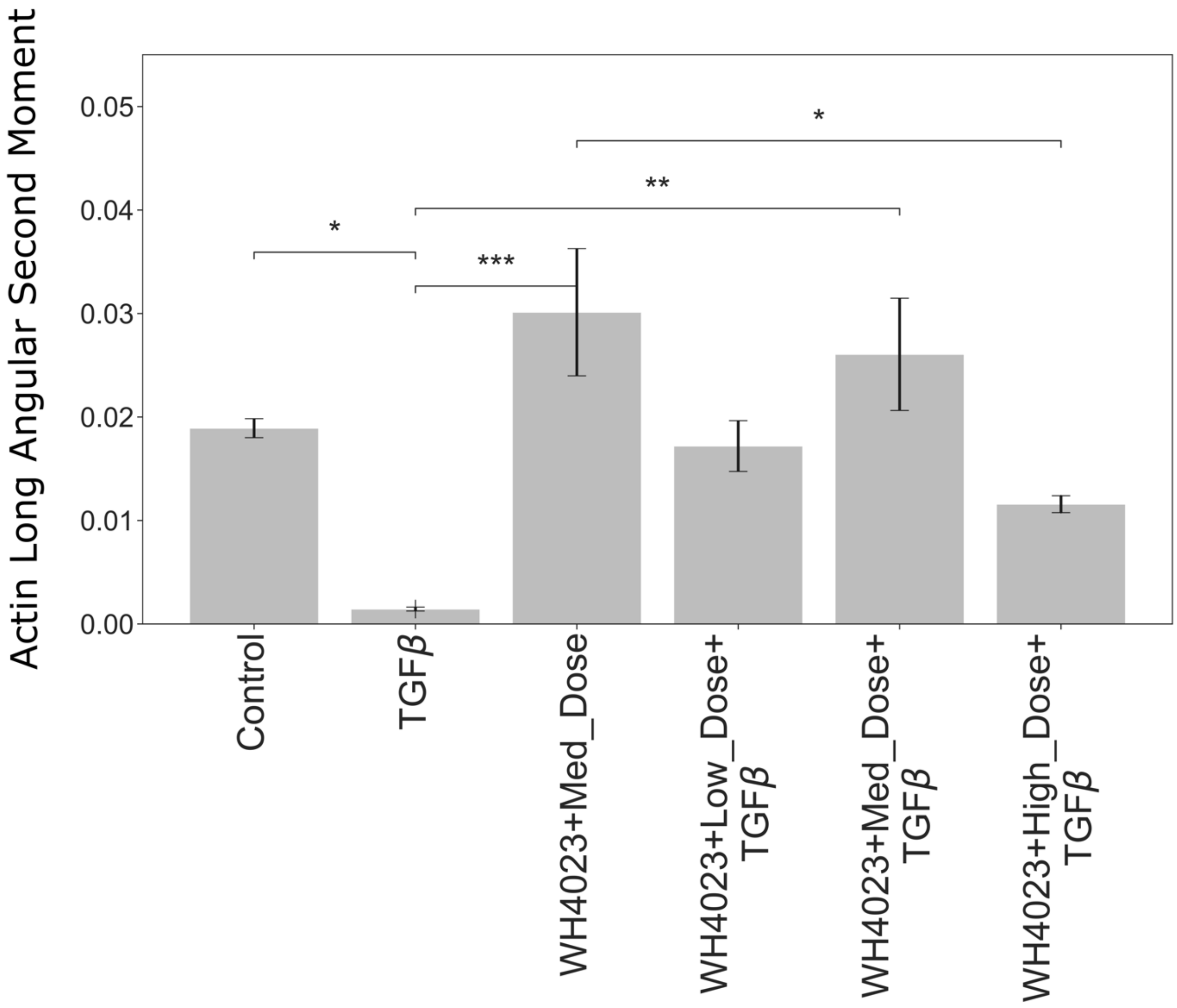


Figure S5: Actin Long Angular Second Moment in response to changing doses of WH. *p-value<=0.05, ANOVA with Tukey’s post-hoc.

| Drug | Modeled Target |
|---|---|
| Anakinra | IL1R1 inhibitor |
| Valsartan | AGIIR inhibitor |
| Defactinib | FAK inhibitor |
| WH-4-023 | Src inhibitor |
| Glutathione | ROS inhibitor |
| CW-HM12 | Integrin inhibitor |
| Salbutamol | BAR2 agonist |
| Marimistat | MMP inhibitor |
| Fasudil | Rock inhibitor |
| SB203580 | p38 inhibitor |
| Pirfenidone | Furin inhibitor |
| Brain natriuretic peptide (BNP) | NPRA agonist |
| Valsartan + BNP | AGIIR inhibitor, NPRA agonist |

Table S 1: Drugs and corresponding targets in the fibroblast network model.

| Feature | Measurement Quantified |
| --- | --- |
| 1) Collagen Integrated Intensity | Total Procollagen I Protein Expression in Cell |
| 2) aSMA Integrated Intensity | Total aSMA Protein Expression in Cell |
| 3) Actin Integrated Intensity | Total Assembled F-actin in Cell |
| 4) Collagen Minimum Intensity | Minimum Procollagen I Protein Expression in Cell |
| 5) Collagen Mean Intensity | Mean Procollagen I Protein Expression in Cell |
| 6) Cell Area | Segmented Cell Size |
| 7) aSMA Long Correlation | Organization of aSMA in Cell, Measured At Long Scale Length |
| 8) Actin Long Angular Second Moment | F-actin Uniformity, Measured At Long Scale Length |
| 9) Collagen Lower Quartile Intensity | Lower Quartile Procollagen I Protein Expression in Cell |
| 10) Actin Long Inverse Difference Moment | Inverse of F-actin Contrast, Measured At Long Scale Length |
| 11) aSMA Long Angular Second Moment | aSMA Uniformity, Measured At Long Scale Length |
| 12) Actin Medium Sum Entropy | F-actin Randomness, Measured At Medium Scale Length |
| 13) aSMA Long Sum Average | Average Normalized aSMA, Measured At Long Scale Length |
| 14) aSMA Medium Sum Entropy | aSMA Randomness, Measured At Medium Scale Length |
| 15) aSMA Long Inverse Difference Moment | Inverse of aSMA Contrast, Measured At Long Scale Length |
| 16) aSMA Long Information Measure 2 | An additional measure of aSMA Information, Measured At Long Scale Length |
| 17) Actin Short Information Measure 1 | A measure of F-actin Information, Measured At Short Scale Length |
| 18) Actin Medium Information Measure 1 | A measure of F-actin Information, Measured At Medium Scale Length |

Table S 2: Retained feature set and measurement quantified for each cell feature.
