## Supplemental Methods for "Logic-based mechanistic machine learning on high-content images reveals how drugs differentially regulate cardiac fibroblasts"

### Construction of the LogiMML model

The LogiMML model was constructed as a two-layer model: first a logic-based differential equation model and then a ridge-regression model.

### Logic-based differential equation model

The first layer is the logic-based fibroblast cell signaling network model. This model is constructed using ordinary differential equations that represent relationships between nodes in key cell signaling pathways. This model uses normalized-Hill equations to represent activation and inhibition of nodes. Differential equations for the model are constructed as follows.

For node C activated by node A and inhibited by node B:

$$\frac{dC}{dt}=\frac{1}{\tau_{C}}(W_{AC}f_{act}\left( A \right)\cdot W_{BC}f_{inhib}\left( B \right)\cdot C_{MAX}-C)$$

Where τ is the time constant for a species, W is the reaction weight between two nodes, bounded in the range 0≤W≤1. Y_MAX_ (0≤ Y_MAX_ ≤1) is the maximum fractional activation of a species. Activation and inhibition functions are represented as follows:

$$f_{act}\left( X \right)=\frac{{BX}^{n}}{K^{n}+X^{n}}; f_{inhib}\left( X \right)=1-\frac{{BX}^{n}}{K^{n}+X^{n}}$$

$$B=\frac{{{EC}_{50}}^{n}-1}{{{2EC}_{50}}^{n}-1};K={(B-1)}^{1/n}$$

Sensitivity analysis was determined as follows, where S_ij_ is the sensitivity of species ‘i’ to a change in species ‘j’:

$$S_{ij}=(\frac{\Delta Y_{i}}{\Delta P_{j}})\cdot(\frac{\Delta Y_{o,i}}{\Delta P_{o,j}})$$

Nodes were perturbed for sensitivity analysis by setting Y_0_(Node)=1 and Tau(Node) =1000000. Cytokine effects were simulated in this layer of the model, by setting inputs weights to 0.25 for baseline cytokines, and 0.6 for high cytokine inputs (i.e. ‘High TGFB Context’), and the predicted node values under each simulated experimental condition were retained for use in the second layer of the LogiMML model.

The first layer of the LogiMML model is an array of predicted node expression values under a set of simulated experimental conditions.

### Ridge regression model mapping network state to fibroblast phenotypes

Mapping all network nodes to a given fibroblast phenotype would be highly underdetermined. Therefore, we first reduced the dimensionality of netwrok model predictions by averaging network nodes within clusters. Cluster labels are assigned to nodes using k-means clustering with k=11. Where X_cluster_ is the X value for a given cluster, and Ynode_i_ represents normalized activity level of a given node in that cluster:

$$X_{cluster}=mean\left( {Ynode}_{i}\ldots{Ynode}_{n} \right)$$

The mean node values for these 11 clusters are set as the X inputs to the second layer of the LogiMML model, the Ridge regression layer. X_cluster_ values were then used to train the Ridge regression model layer and determine the following model equation:

$$Y_{feature}={\beta_{0}+X}_{1}\beta_{1}+X_{2}\beta_{2}\ldots X_{n}\beta_{n}$$

Where Y_feature_ is the measured value of a given cell feature, X is the predicted mean expression of a cluster, $\beta$ is the beta coefficient for the corresponding X term, or the intercept term in the case of $\beta$_0_ and n is the total number of clusters (n=11).

In this ridge-regression layer, the 11 averaged node values from a given simulated condition of interest (i.e. ‘TGFB+Fasudil’) are fed into the model as X values. The Y value for training the model is the experimentally determined value for a feature of interest (i.e. Integrated Collagen Intensity) under the corresponding drug treatment of interest. The Ridge regression is then trained on 56 sets of X and Y inputs corresponding to 56 experimental conditions (negative control, cytokine treatments, and maximum dose for drug and ‘drug+cytokine’ groups). The model is trained using the ‘linear_model.Ridge’ function from the ‘sklearn’ Python package v1.3.2. Ridge regression was chosen because it gave a broader range of predictions involving multiple candidate pathways, whereas elastic net and LASSO regression methods typically focused the weights towards a single node.

### Using the combined LogiMML model (network + ridge regression) to simulate fibroblast phenotypes in response to new perturbations

The trained model is then run using new X data from simulations that correspond to hypothetical experimental conditions (i.e. ‘TGFB+Fasudil with a Rho knockout’). As the inputs to the second layer of the model are normalized values, the outputs, or Y values from this model are also normalized. Full code to reproduce and implement this mode is available at https://github.com/andersnelson/Logic-based_MML.
